## Supplementary filr for "Rad52 Acts as an Assembly Chaperone to Form and Stabilize Rad51 Filaments Through a Large C-Terminus 85-Residue Segment"

*co-corresponding

[1]Université Paris Cité, Inserm, CEA, Stabilité Génétique Cellules Souches et Radiations, LRGM/iRCM/IBFJ, F-92260 Fontenay-aux-Roses, France.
[2]Université Paris-Saclay, Inserm, CEA, Stabilité Génétique Cellules Souches et Radiations, LRGM/iRCM/IBFJ, F-92260 Fontenay-aux-Roses, France.
[3]Nuclear Dynamics, CNRS UMR 3664, Institut Curie, PSL Research University, Sorbonne Université, Paris 75005, France.
[4]Institute Joliot, Commissariat à l'énergie Atomique (CEA), Direction de la Recherche Fondamentale (DRF), F91191 Gif-sur-Yvette, France
[5]Institute for Integrative Biology of the Cell (I2BC), CEA, CNRS, Univ. Paris-Sud, Université Paris-Saclay, 91198, Gif-sur-Yvette cedex, France
[6]Université Paris Cité, Inserm, CEA, Stabilité Génétique Cellules Souches et Radiations, CIGEx/iRCM/IBFJ, F-92260 Fontenay-aux-Roses, France.
[7]Université Paris-Saclay, Inserm, CEA, Stabilité Génétique Cellules Souches et Radiations, CIGEx/iRCM/IBFJ, F-92260 Fontenay-aux-Roses, France.
[8]Synchrotron SOLEIL, HelioBio group, l'Orme des Merisiers, Départementale 128, 91190 Saint-Aubin, France.

**Supplementary Figures**

**S1 to S7**

**Supplementary Tables**

**Tables S1 to S2**

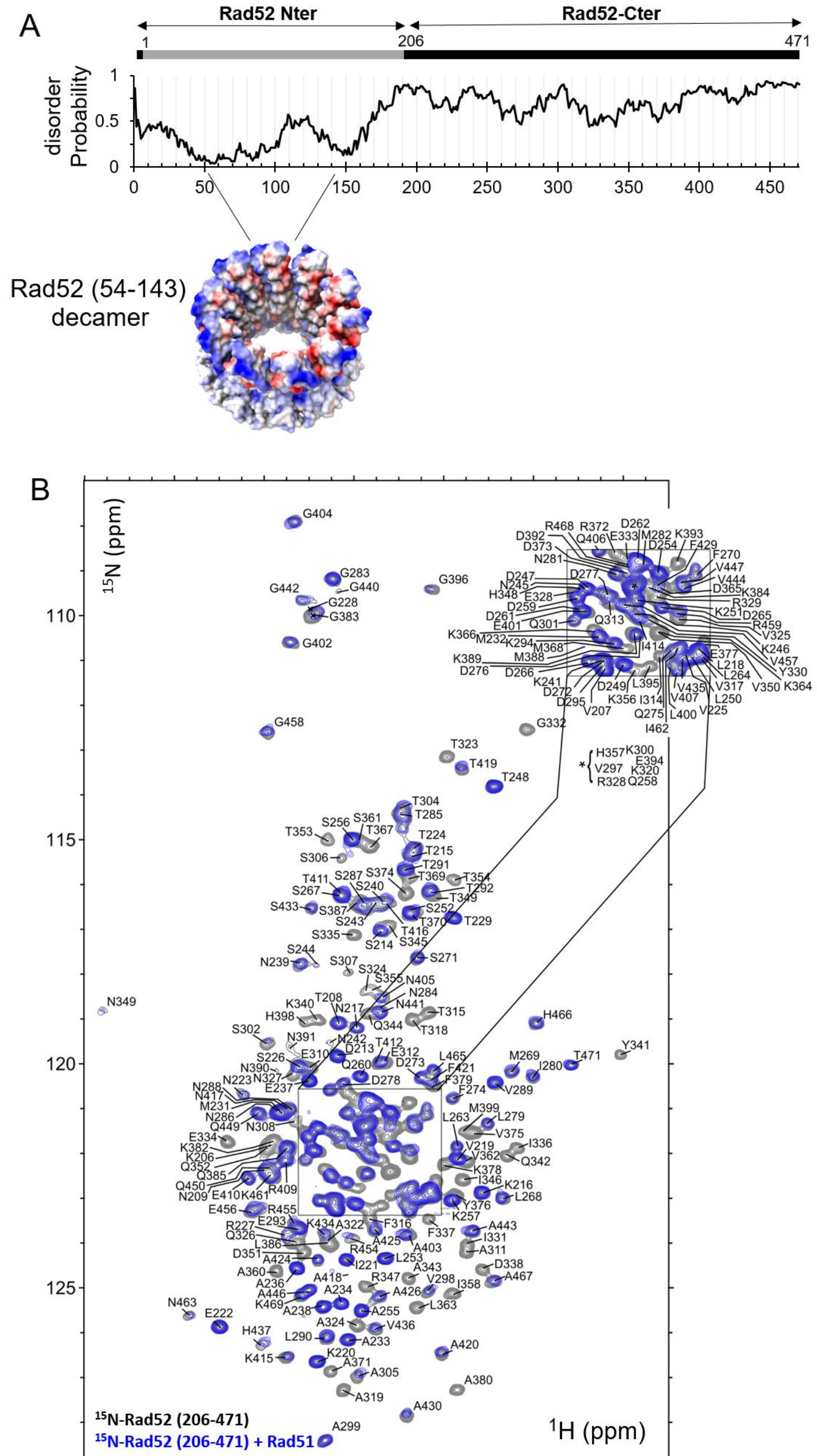
A
Rad52 Nter
Rad52-Cter
1
206
471
disorder Probability
1
0.5
0
0
50
100
150
200
250
300
350
400
450
Rad52 (54-143) decamer
B
$^{15}$N (ppm)
110
115
120
125
$^{1}$H (ppm)
9.0
8.8
8.6
8.4
8.2
8.0
7.8
$^{15}$N-Rad52 (206-471)
$^{15}$N-Rad52 (206-471) + Rad51

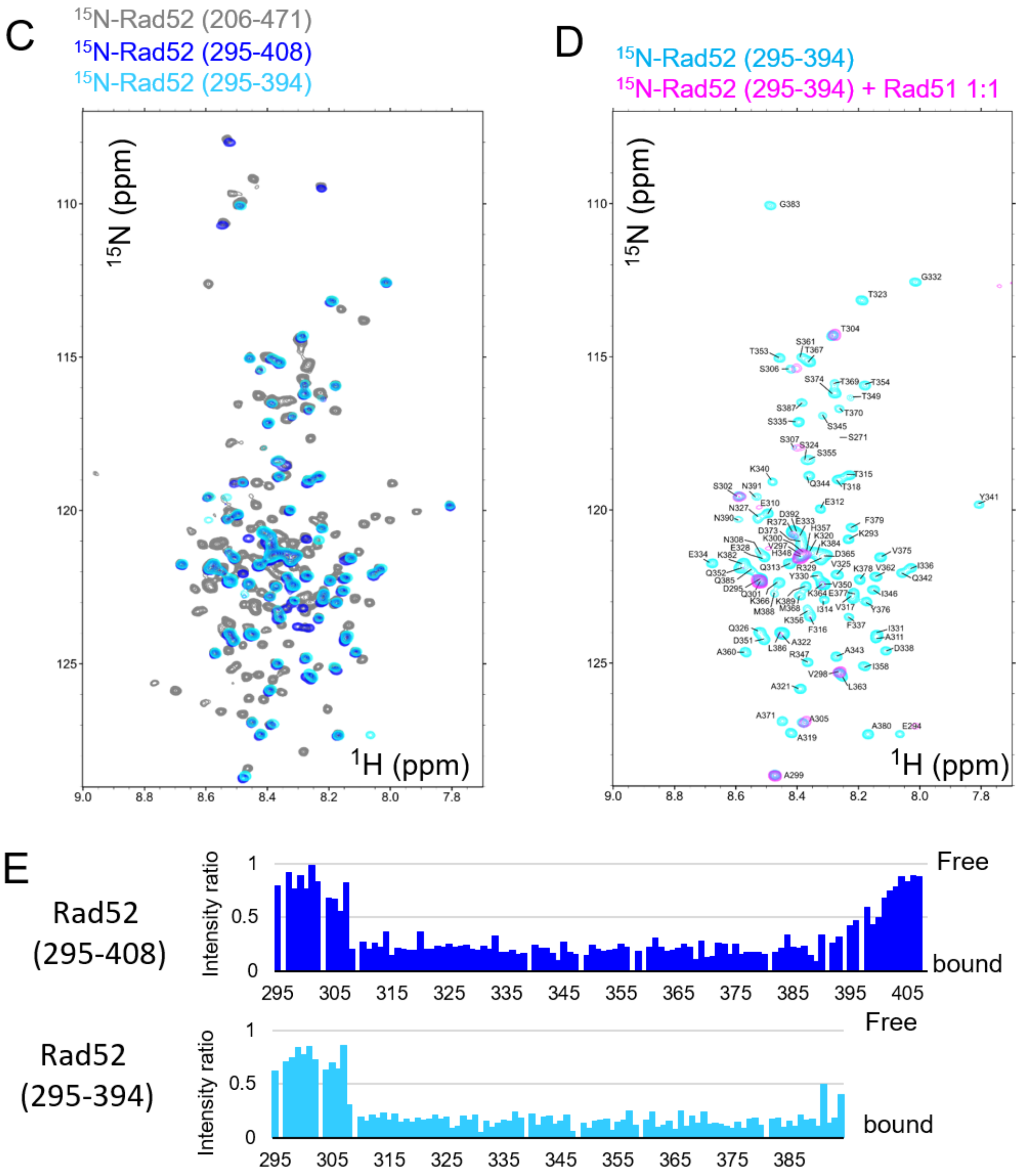

**Supplementary Figure 1: Rad52 C-terminus is disordered and interacts with Rad51 with a central region of 85 residues**

**A:** General organisation of the N-terminal and C-terminal domains of Rad52 and the disorder probability for the full Rad52 sequence. The experimental decameric structure of the N-terminal region of yeast Rad52 (PDB:°8G3G) is shown as a surface with electrostatic colouring from blue for positive changes to red for negative charges **B:** $^{1}H$-$^{15}N$ SOFAST-HMQC spectra of the uniformly $^{15}N$ labelled Rad52-Cter domain (206-471) alone in grey and, after addition of equimolar amount of unlabeled Rad51 in blue. The full assignment of residues is indicated. **C:** $^{1}H$-$^{15}N$ SOFAST-HMQC spectra of uniformly $^{15}N$ labelled Rad52-Cter domain (206-471) in grey, (295-408) in blue and (295-394) in cyan. **D:** $^{1}H$-$^{15}N$ SOFAST-HMQC spectra of the uniformly $^{15}N$ labelled Rad52-Cter domain (295-394) alone in cyan and, after addition of equimolar amount of unlabeled Rad51 in magenta. The assignment of residues is indicated. **E:** Mapping of the interaction between Rad52-Cter domains (295-408 in the upper panel) or (295-394 in the lower pane) with Rad51, using the intensities ratio (I/I0), where I and I0 are the intensity of the signals $^{1}H$-$^{15}N$ SOFAST-HMQC spectra before and after addition of Rad51, respectively.

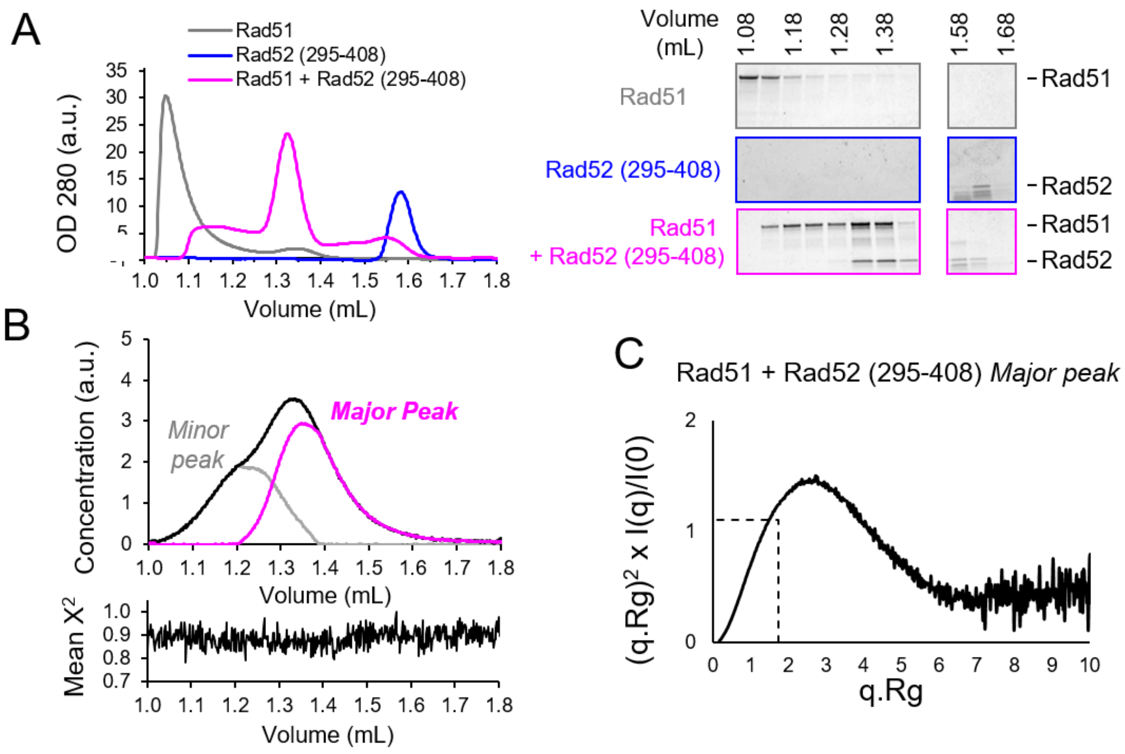

**Supplementary Figure 2: Rad52 C-ter is disordered and interacts with Rad51 with central region of 85 residues**

**A:** Left panel: SEC profile analysis of Rad51, Rad52 (295-408) and Rad51+Rad52 (295-408) in a 1:1 ratio. Right panel: SDS-PAGE analysis of fraction from SEC profile in left panel revealed with coomassie blue. **B:** Deconvolution of the SEC-SAXS curve, upper panel, area-normalized concentration profiles for each component. With the major peak shown in magenta and the minor peak in grey. Lower panel: Mean $\chi^2$ values of the fit of the deconvolution and the original data. **C:** Dimensionless Kratky plot of the SAXS experiment of Rad51 + Rad52 (295-408) major peak after deconvolution. The expected maximum position for a fully globular protein is indicated by dashed black lines.

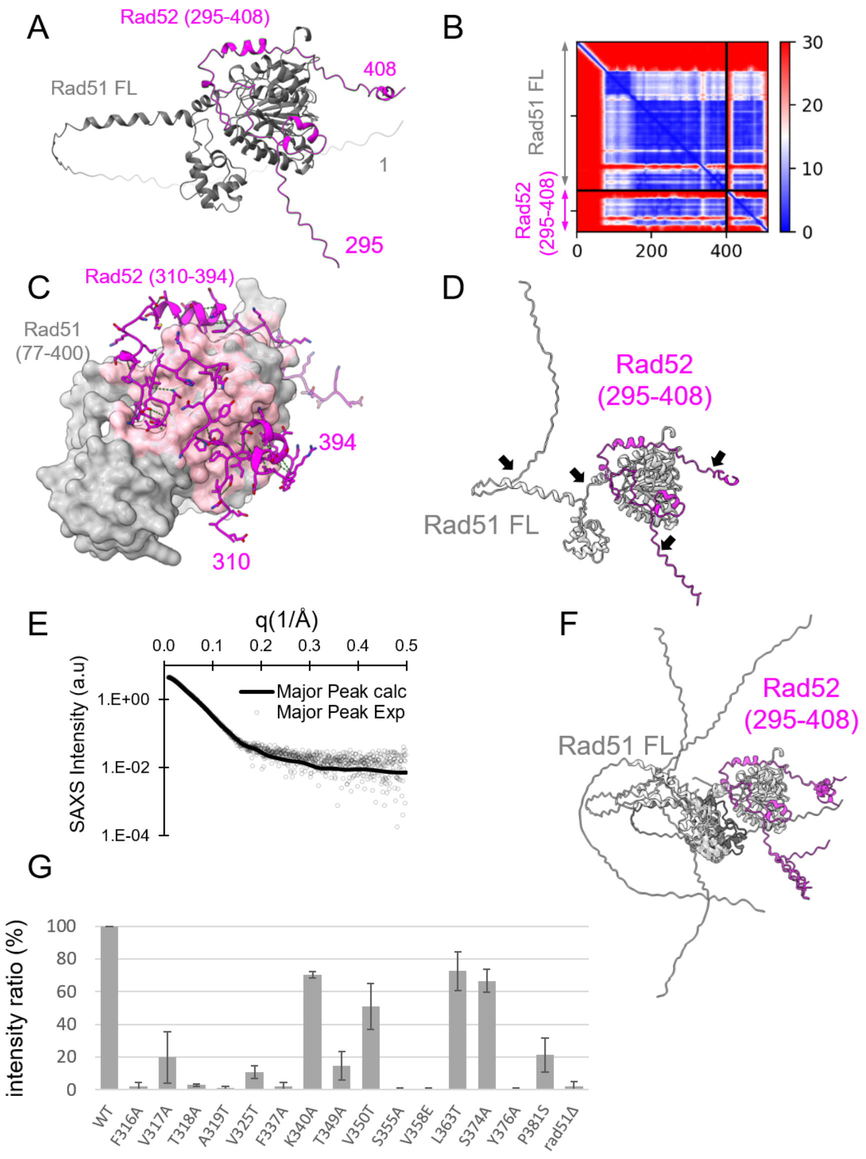

**Supplementary Figure 3: Rad52 C-ter is disordered and interacts with Rad51 with central region of 85 residues**

**A:** AF2 model of the complex between Rad51 + Rad52 (295-408). Both proteins are shown as cartoons, with Rad51 in grey and Rad52 in magenta **B:** Predicted Alignment Error plot (PAE) calculated by AlphaFold2 for the best model of Rad51+Rad52 (295-408) presented in panel A. **C:** AF2 model of the complex between Rad51 (77-400) + Rad52 (310-394). Rad51 is displayed as a surface, with residues in contact with Rad52 colored pink and other residues in grey. Rad52 is shown as a magenta cartoon, with visible side chains. **D:** Ribbon representation of the best model generated by the Dadimodo software [1]. Both proteins are shown as cartoons, with Rad51 in grey and Rad52 in magenta. The position of the last residue of the N-terminal tail of Rad51, and the hinge between the N-terminal four-helix domain and the C-terminal domain are indicated with black arrows. the positions of residues 310 and 394, which delimit the two extremities considered flexible in Rad52, are also indicated with black arrows **E:** Experimental and fitted SAXS profile intensity (I) as a function of the momentum transfer (q) for Rad51 + Rad52 (295-408) major peak after deconvolution (black circles) and for the best model generated by the Dadimodo Software (continuous black line) shown in panel D. **F:** Overlay of the 8 best models generated by the Dadimodo software in light grey and the initial model generated by AlphaFold in dark grey. Rad52 is shown in magenta **G:** Intensity ratio between mutant and WT Rad52 co-immunoprecipitated with Rad51. Data are presented as the mean ± SEM for at least 2 two experiments.

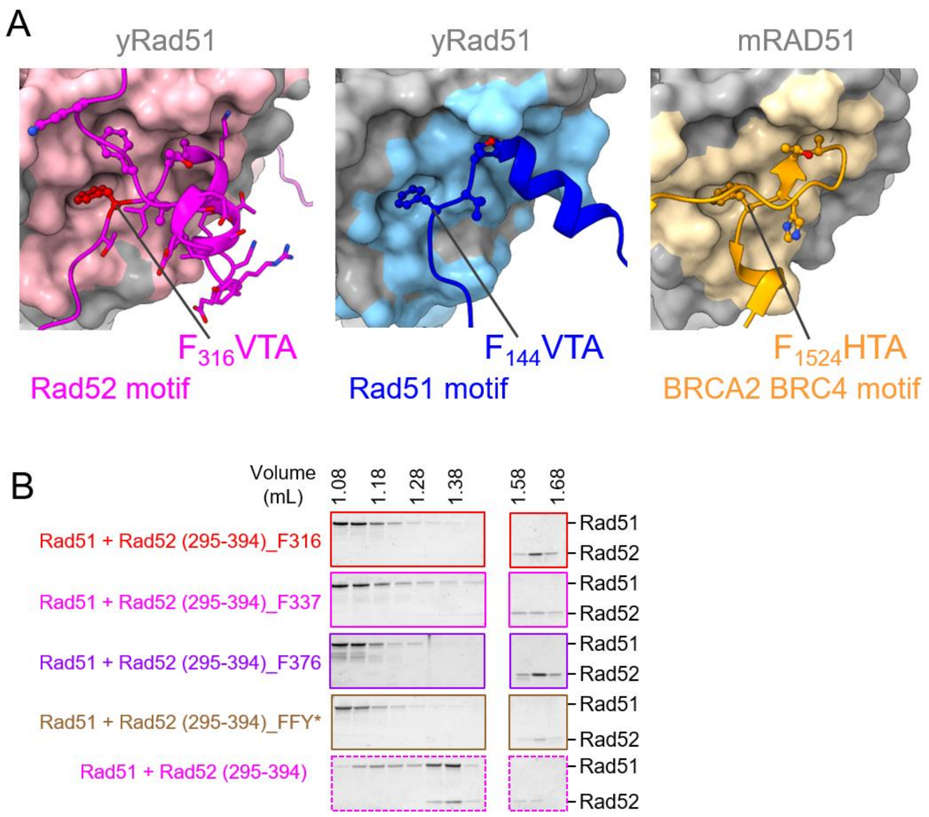

**Supplementary Figure 4: Rad52 (310–394) competes with Rad51 oligomers through the cooperative binding of several anchors**

**A:** Zoomed-in view of the region of Rad51 interacting with FxxA motif. Left panel AF2 model of the complex between Rad51 and Rad52. Rad51 is displayed as a surface, with residues in contact with Rad52 colored pink and other residues in grey. Rad52 is shown as a magenta cartoon, with visible side chains. Middle panel, interaction of two Rad51 monomers assembled in a multimer (PDB code 1SZP). One Rad51 monomer is shown as a surface, with residues in contact with the second Rad51 colored blue and other residues in grey. The second monomer, containing the FVTA motif, is displayed as a blue cartoon, with visible side chains for the FVTA motif. Right panel Structure of mouse Rad51 interacting with the FHTA BRC4 motif of BRCA2 (PDB code 1N0W). Rad51 is shown as a surface, with residues in contact with BRCA2 colored orange. BRCA2 is represented as an orange cartoon, with visible side chains for the FHTA motif. **B:** SDS-PAGE analysis of fractions from SEC profiles in **Figure 4D,** stained with coomassie blue.

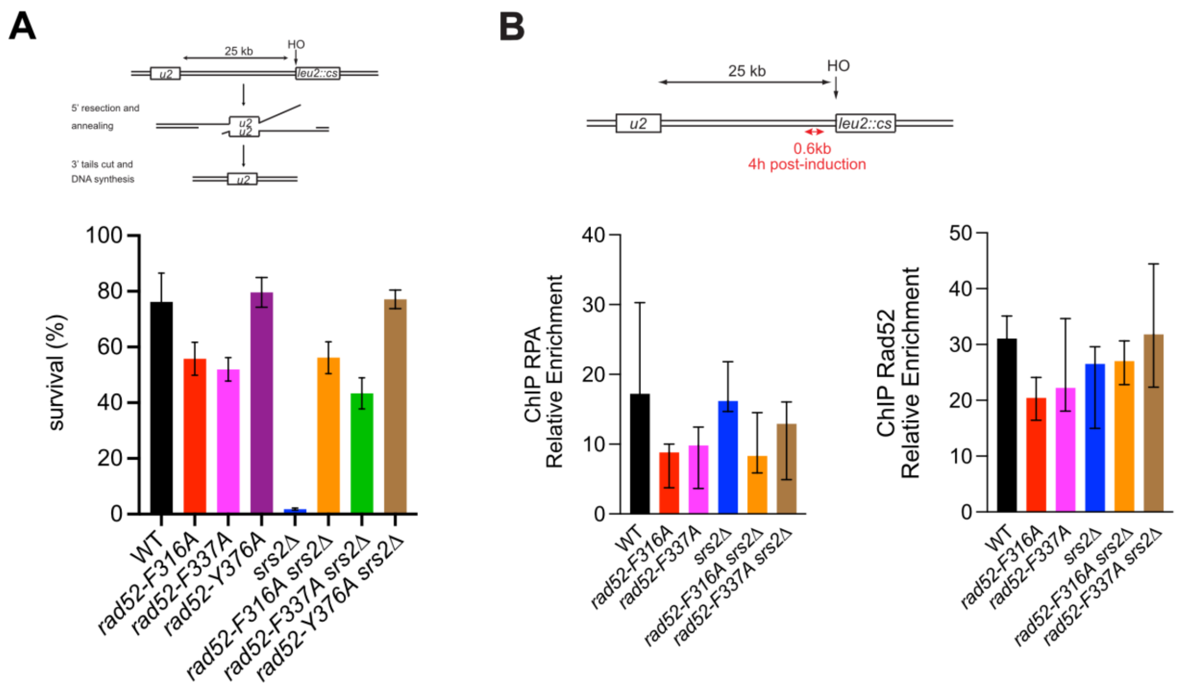

**Supplementary Figure 5**

**A:** Upper panel: schematic of the different steps of an HO-induced DSB repair by SSA. Cell survival after HO-induced DSB formation in a SSA repair system. Lower panel: data are presented as the mean ± SEM of at least three independent experiments. Already published data obtained with *rad52-Y376A* are shown for reference. **B:** Upper panel: schematic of the HO-induced SSA repair system is shown. Lower panel: ChIP was used to assess RPA and Rad52 relative enrichment at 0.6 kb from the DSB site (red) 4 hours after HO induction. Data are presented as the median and minimum and maximum values of at least three independent experiments; None of the results are statistically significant (two-tailed unpaired t-test).

| **Impact of *rad52* mutations** | | | |
|---|---|---|---|
| Conditions | p-value filament length | p-value filament intensity | p-value Foci intensity |
| *WT vs F337A* | n.a. | n.a. | 4.07 e-6 *** |
| *WT vs Y376A* | 0.569 | 0.402 | 1.95 e-5 *** |
| *WT vs F316A* | n.a. | n.a. | 1.63 e-6 *** |

| **Impact of *SRS2* deletion on WT and *rad52* mutant strains** | | | |
|---|---|---|---|
| Conditions | p-value filament length | p-value filament intensity | p-value Foci intensity |
| *WT vs srs2Δ* | 0.590 | 1.02 e-8 *** | 0.0171* |
| *F337A vs F337A srs2Δ* | n.a. | n.a. | 2.10 e-12 *** |
| *Y376A vs Y376A srs2Δ* | 0.865 | 0.164 | 5.89 e-9 *** |
| *F316A vs F316A, srs2Δ* | n.a. | n.a. | 6.08 e-16 *** |

| **Impact of *rad52* mutations on *SRS2* deleted strains** | | | |
|---|---|---|---|
| Conditions | p-value filament length | p-value filament intensity | p-value Foci intensity |
| *srs2Δ vs F337A srs2Δ* | 0.0363* | 7.28 e-12 *** | 4.20 e-5 *** |
| *srs2vs Y376A srs2Δ* | 0.0150* | 8.00 e-13 *** | 0.0101* |
| *srs2 vs F316A srs2Δ* | 0.0336* | 7.14 e-5 *** | 7.05 e-7 *** |

| ***srs2Δ rad52 mutants vs* WT** | | | |
|---|---|---|---|
| Conditions | p-value filament length | p-value filament intensity | p-value Foci intensity |
| *WT vs F337A srs2Δ* | 3.41 e-6 *** | 0.210 | 0.275 |
| *WT vs Y376A srs2Δ* | 4.56 e-7 *** | 1 | 1 |
| *WT vs F316A srs2Δ* | 0.654 | 5.30 e-8 *** | 4.56 e-7 *** |

**Supplementary Figure 6: Statistical test corresponding to Figure 6.** Logistic regression with binomial distribution (see materials and methods; ***P < 0.001, **P < 0.01, *P < 0.05).

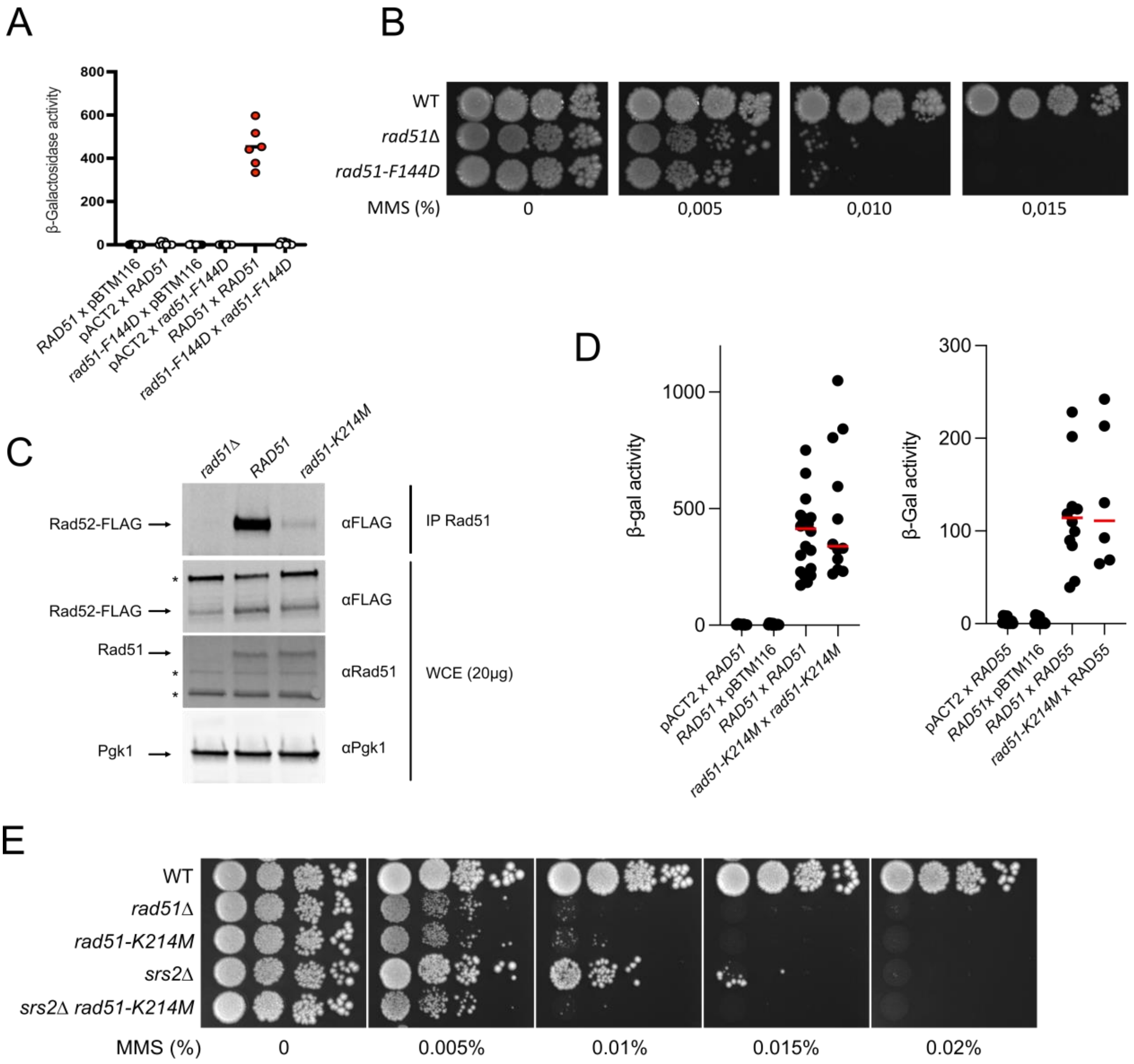

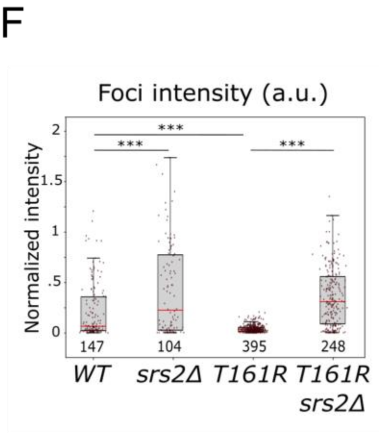

G

| Conditions | p-values Filament Intensity | p-values Filament length | p-values Foci Intensity |
|---|---|---|---|
| *wt vs srs2* | 9.14 e-12 *** | 0.847 | 5.93 e-5 *** |
| *wt vs T161R* | 1.26 e-3 * | 0.847 | 3.12 e-11 *** |
| *wt vs srs2 T161R* | 3.38 e-3 * | 1.79 e-8 *** | 3.47 e-5 *** |
| *T161R vs T161R srs2* | 8.85 e-4 ** | 0.847 | 9.13 e-21 *** |
| *srs2 vs T161R srs2* | 4.31 e-5 *** | 8.08 e-9 *** | 0.0673 |

**Supplementary Figure 7. Rad51-F144D is defective in Rad51-Rad51 interaction and Rad51-K214M is highly defective in HR.**

**A**: Y2H analysis of Rad51-F144D interaction with itself. Semi-quantitative measurement of β-galactosidase activity in independent colonies issued from co-transformation of plasmids bearing the LexA-activating domain (pACT2) and the GAL4 binding domain (pBTM) either empty or bearing fusion proteins. Note that in addition of F144D, *RAD51* is also mutated for E221K, which suppresses the toxicity of Rad51 overexpression (see Materials and Methods). **B**: Serial 10-fold dilutions of haploid strains with the indicated genotypes were spotted onto rich medium (YPD) containing different MMS concentrations. **C:** Co-IP experiments showing the loss of interaction between Rad51-K214M and Rad52. Rad51 was immunoprecipitated with a poly-clonal anti-Rad51 antibody (αRad51). The presence of Rad51 in the immunoprecipitated fraction (IP) cannot be detected because it migrates at the same level as the anti-Rad51 IgG used for the immunoprecipitation. However, the absence of Rad52-FLAG in the *rad51*Δ immunoprecipitate confirmed that the Rad52-FLAG signal observed is related to the Rad52–Rad51 interaction. Western blot analysis of Rad52-FLAG, Rad51 and Pgk1 in whole cell extracts (WCE) reveals that the amount of proteins is comparable in each mutant strains. * Unspecific bands. **D:** Y2H analysis of Rad51-K214M interaction with itself and with Rad55-Rad57. Semi-quantitative measurement of β-galactosidase activity in independent colonies issued from co-transformation of plasmids bearing the LexA-activating domain (pACT2) and the GAL4 binding domain (pBTM) either empty or bearing fusion proteins. Note that in addition of K214M, *RAD51* is also mutated for E221K, which suppresses the toxicity of Rad51 overexpression (see Materials and Methods). **E:** Serial 10-fold dilutions of haploid strains with the indicated genotypes were spotted onto rich medium (YPD) containing different MMS concentrations. **F**: Comparison of Rad51 foci intensities in WT and mutant strains as indicated. The number of structures analyzed is indicated below each box. Intensities are normalized to the mean intensity of WT Rad51 filaments. On each box, the central mark indicates the median, and the bottom and top edges of the box indicate the 25th and 75th percentiles, respectively. The whiskers extent corresponds to the adjacent value, which is the most extreme data value not considered an outlier (above 75th percentile + 1.5 times interquartile range or below 25th percentile - 1.5 times interquartile range). **G**: Statistical test corresponding to Figure 7F. Logistic regression with binomial distribution (see materials and methods; ***P < 0.001, **P < 0.01, *P < 0.05).

### Supplementary Table 1: Experimental information and modelling of SAXS data

| **Sample details** | **Rad51 + Rad52 (295-308)** |
|---|---|
| Organism | *S. cerevisiae* |
| Source (catalogue No. or reference) | Recombinant proteins (See Methods) |
| UniProt sequence ID (residues in construct) | P25454(FL 1-400) P06778 (295-408) |
| Extinction coefficient [$A_{280}$, 0.1%(*w/v*)] | 0.295 |
| M from chemical composition (Da) | 55 584 |
| SEC–SAXS column | S200 3.2/300 Increase |
| Loading concentration (mg $ml^{-1}$) | 1.3 |
| Injection volume (µl) | 50 |
| Flow rate (ml $min^{-1}$) | 0.05 |
| Solvent (solvent blanks taken from SEC flowthrough prior to elution of protein) | Tris 20 mM, NaCl 100 mM, pH 8 |
| **SAXS data-collection parameters.** | |
| Instrument/data processing | BioSAXS on the SWING beamline at Synchrotron SOLEIL(Thureau et al. 2021) |
| Wavelength (Å) | 1.0332 |
| Beam size (µm) | 500x200 |
| Camera length (m) | 2.00 |
| q measurement range ($Å^{-1}$); $q = 4\pi\sin(\theta)/\lambda$ (2θ: scattering angel & λ the x-ray wavelength) | 0.00365–0.5538 |
| Absolute scaling method | Comparison with scattering from 1 mm pure $H_2O$ |
| Normalization | To transmitted intensity by beam-stop counter |
| Monitoring for radiation damage | data frame-by-frame comparison |
| Exposure time | Continuous 1 s data-frame measurements of SEC elution |
| Sample configuration | SEC–SAXS with thermalized quartz capillary (ID 1.5mm) |
| Sample temperature (°C) | 20 |
| **Software employed for SAXS data reduction, analysis and interpretation** | |
| SAXS data reduction | I(q) versus q, buffer subtraction & frames selection using Foxtrot 3.10 [a] |
| Extinction coefficient estimate | ProtParam [2] |
| Basic analyses: Guinier, *P(r)*, MW | PRIMUSqt from ATSAS 3.2.1 [3] |
| Atomic structure modelling | Dadimodo [1] (https://dadimodo.synchrotron-soleil.fr/) |
| Missing sequence modelling | AlphaFold2 [4] |
| Three-dimensional graphic model representations | PyMOL v.2.00 |
| **Structural parameters** | |
| Guinier analysis | |
| I(0) ($cm^{-1}$) | 4.75 ± 5E-3 |
| $R_g$ (Å) | 39.36 ± 0.06 |
| $q_{min}$ ($Å^{-1}$) | 0.0114 |
| $qR_g$ max (q min = 0.0066 $Å^{-1}$) | 1.33 |
| Coefficient of correlation, $R^2$ | 0.998 |
| M from Vc (ratio to predicted) | 68 300 (0.81) |
| P(r) analysis | |
| I(0) ($cm^{-1}$) | 4.86 ± 9E-3 |
| $R_g$ (Å) | 43.0 ± 0.19 |
| $d_{max}$ (Å) | 170 |
| q range ($Å^{-1}$) | 0.011 to 0.554 |
| total estimate from GNOM | 0.724 |
| **Atomistic modelling.** | |
| Crystal structures | |
| q range for all modelling | 0.01003–0.49998 |
| PepsiSAXS (r0 fixed) | |
| $\chi^2$ | 2.69 |
| Vol ($Å^3$), R0 (Å), Dro (e $Å^{-3}$) | 68043, 1.615, 0.00334 |
| Dadimodo (https://dadimodo.synchrotron-soleil.fr/) | |
| Starting structures | From AlphaFold2 |
| Rigid bodies | body1 = A: 81-141<br>body2 = A: 156-400, B: 17-96 |
| No. of generated structures | 9 |
| $\chi^2$ range from PepsiSAXS | 2.50 - 2.94 |

[a] (https://www.synchrotron-soleil.fr/en/beamlines/swing#paragraphes_menu_left-block-7)

### Supplementary Table 2. *S. cerevisiae* strains.

| Strain | Genotype | Figure | Source | Strain background |
|---|---|---|---|---|
| FF18733 | *MAT*a *leu2-3, 112 trp1-289 ura3-52 lys1-1 his7-2* | Fig6-Ex1B | F. Fabre | FF18733 |
| ECS2917 | *rad52::URA3 YCplac111* | Fig. 3D, 3-Ex3, 5A | [5] | |
| ECS2947 | *rad52::URA3 YCplac111-RAD52-6His-3FLAG* | Fig. 3D, 3-Ex3, 5A | [5] | |
| ECS2928 | *rad51::URA3 YCplac111-RAD52-6His-3FLAG* | Fig. 3D, 3-Ex3 | [5] | |
| ECS3033 | *rad52::URA3 YCplac111-rad52-F316A-6His-3FLAG* | Fig. 3D, 3-Ex3, 5A | | |
| ECS3035 | *rad52::URA3 YCplac111-rad52-V317A-6His-3FLAG* | Fig. 3D, 3-Ex3 | | |
| ECS3037 | *rad52::URA3 YCplac111-rad52-T318A-6His-3FLAG* | Fig. 3D, 3-Ex3 | | |
| ECS3039 | *rad52::URA3 YCplac111-rad52-A319T-6His-3FLAG* | Fig. 3D, 3-Ex3 | | |
| ECS3045 | *rad52::URA3 YCplac111-rad52-V325T-6His-3FLAG* | Fig. 3D, 3-Ex3 | | |
| ECS3047 | *rad52::URA3 YCplac111-rad52-I331T-6His-3FLAG* | Fig. 3D, 3-Ex3 | | |
| ECS3051 | *rad52::URA3 YCplac111-rad52-F337A-6His-3FLAG* | Fig. 3D, 3-Ex3, 5A | | |
| ECS3055 | *rad52::URA3 YCplac111-rad52-K340A-6His-3FLAG* | Fig. 3D, 3-Ex3 | | |
| ECS3061 | *rad52::URA3 YCplac111-rad52-V350T-6His-3FLAG* | Fig. 3D, 3-Ex3 | | |
| ECS3063 | *rad52::URA3 YCplac111-rad52-S355A-6His-3FLAG* | Fig. 3D, 3-Ex3 | | |
| ECS3065 | *rad52::URA3 YCplac111-rad52-L363T-6His-3FLAG* | Fig. 3D, 3-Ex3 | | |
| ECS3071 | *rad52::URA3 YCplac111-rad52-P438H-6His-3FLAG* | Fig. 3D, 3-Ex3 | | |
| ECS2987 | *rad52::URA3 YCplac111-rad52-S374A-6His-3FLAG* | Fig. 3D, 3-Ex3 | | |
| ECS2929 | *rad52::URA3 YCplac111-rad52-Y376A-6His-3FLAG* | Fig. 3D, 3-Ex3, 5A | [6] | |
| ECS2930 | *rad52::URA3 YCplac111-rad52-K378A-6His-3FLAG* | Fig. 3D, 3-Ex3 | [6] | |
| ECS2931 | *rad52::URA3 YCplac111-rad52-P381S-6His-3FLAG* | Fig. 3D, 3-Ex3 | [6] | |
| ECS2932 | *rad52::URA3 YCplac111-rad52-G383A-6His-3FLAG* | Fig. 3D, 3-Ex3 | [6] | |
| ECS3772 | *rad52::KanMX rad51 ::KanMX YCplac111-RAD51 YCplac33* | | | |
| ECS3800 | *rad52::KanMX rad51 ::KanMX YCplac111-RAD51 YCplac33-rad52-K340E-6His-3FLAG* | Fig. 3D, 3-Ex3 | | |
| ECS3802 | *rad52::KanMX rad51 ::KanMX YCplac111-RAD51 YCplac33-rad52-T349A-6His-3FLAG* | Fig. 3D, 3-Ex3 | | |
| ECS3796 | *rad52::KanMX rad51 ::KanMX YCplac111-RAD51 YCplac33-rad52-S355A-6His-3FLAG* | Fig. 3D, 3-Ex3 | | |
| ECS3804 | *rad52::KanMX rad51 ::KanMX YCplac111-RAD51 YCplac33-rad52-V358E-6His-3FLAG* | Fig. 3D, 3-Ex3 | | |
| ECS3710 | *rad52 ::KanMX YCplac33* | Fig. 3E | | |
| ECS3712 | *rad52 ::KanMX YCplac33-RAD52* | Fig. 3E | | |
| ECS3567 | *rad52 ::URA3 YCplac111-rad52-F316A* | Fig. 3E | | |
| ECS3569 | *rad52 ::URA3 YCplac111-rad52-F337A* | Fig. 3E | | |
| ECS3663 | *rad52 ::KanMX YCplac33-rad52-K340E* | Fig. 3E | | |

| | | |
|---|---|---|
| ECS3665 | *rad52 ::KanMX YCplac33-rad52-T349A* | Fig. 3E |
| ECS3671 | *rad52 ::KanMX YCplac33-rad52-S355A* | Fig. 3E |
| ECS3667 | *rad52 ::KanMX YCplac33-rad52-V358E* | Fig. 3E |
| ECS3565 | *rad52 ::URA3 YCplac111-rad52-Y376A* | Fig. 3E |
| ECS3738 | *rad52 ::KanMX srs2 ::NatMX YCplac33-RAD52* | Fig. 3E |
| ECS3577 | *rad52 ::URA3 srs2 ::NatMX YCplac111-rad52-F316A* | Fig. 3E |
| ECS3579 | *rad52 ::URA3 srs2 ::NatMX YCplac111-rad52-F337A* | Fig. 3E |
| ECS3730 | *rad52 ::KanMX srs2 ::NatMX YCplac33-rad52-K340E* | Fig. 3E |
| ECS3732 | *rad52 ::KanMX srs2 ::NatMX YCplac33-rad52-T349A* | Fig. 3E |
| ECS3740 | *rad52 ::KanMX srs2 ::NatMX YCplac33-rad52-S355A* | Fig. 3E |
| ECS3734 | *rad52 ::KanMX srs2 ::NatMX YCplac33-rad52-V358E* | Fig. 3E |
| ECS3575 | *rad52 ::URA3 srs2 ::NatMX YCplac111-rad52-Y376A* | Fig. 3E |

| | | |
|---|---|---|
| ECS3031 | *rad52 ::URA3 srs2 ::NAT YCplac111-RAD52-6His-3FLAG* | Fig. 5A |
| ECS3072 | *rad52 ::URA3 srs2 ::NAT YCplac111-RAD52-F316A-6His-3FLAG* | Fig. 5A |
| ECS3090 | *rad52 ::URA3 srs2 ::NAT YCplac111-RAD52-F337A-6His-3FLAG* | Fig. 5A |
| ECS2937 | *rad52 ::URA3 srs2 ::NAT YCplac111-RAD52-Y376A-6His-3FLAG* | Fig. 5A [2] |

| | | |
|---|---|---|
| ECS3768 | *rad52::KanMX rad51 ::KanMX YCplac111-RAD51 YCplac33-RAD52-6His-3FLAG* | Fig. 6A, 7A |
| ECS3770 | *rad52::KanMX rad51 ::KanMX YCplac111 YCplac33-RAD52-6His-3FLAG* | Fig. 6A, 7A |
| ECS3752 | *rad52::KanMX rad51 ::KanMX YCplac111-rad51-K214M YCplac33-RAD52-6His-3FLAG* | Fig. 6A |
| ECS3764 | *rad52::KanMX rad51 ::KanMX YCplac111-rad51-T161R YCplac33-RAD52-6His-3FLAG* | Fig. 7A |

| | | |
|---|---|---|
| ECS3555 | *rad51 ::NatMX YCplac111-RAD51* | Fig. 6C, 7B, 7C |
| ECS3557 | *rad51 ::NatMX YCplac111* | Fig. 6C, 7B |
| ECS3541 | *rad51 ::NatMX YCplac111-RAD51-K214M* | Fig. 6C |
| ECS3559 | *rad51 ::URA3 YCplac111-RAD51 srs2 ::NAT* | Fig. 6C, 7B, 7C |
| ECS3587 | *rad51 ::URA3 YCplac111-RAD51-K214M srs2 ::NAT* | Fig. 6C |
| ECS3553 | *rad51 ::NatMX YCplac111-RAD51-T161R* | Fig. 7B, 7C |
| ECS3599 | *rad51 ::URA3 YCplac111-RAD51-T161R srs2 ::NAT* | Fig. 7B, 7C |

| | | |
|---|---|---|
| ECS3930 | *rad51-F144D* | Fig6-Ex1B |
| TLM297 | *Rad51 ::NATMX arg4-Bg* | Fig6-Ex1B |

| | | | | |
|---|---|---|---|---|
| tGI354 | *MAT***a** *hml::ADE1 MATalpha hmr::ADE1 arg5,6::MATa-inc::HPH1 ade3::GAL::HO* | Fig. 5B | J. Haber | JKM146 |
| ECS3622 | *rad52-F316A* | Fig. 5B | | |
| ECS3625 | *rad52-F337A* | Fig. 5B | | |
| TLM208 | *Rad52-Y376A* | Fig. 5B | [6] | |
| yAT3880 | *MAT***a** *can1-100 his3-11,15 leu2-3,112 ura3Δ::KanMX RAD5 lys2::ura3-ISceIcutsite (loxP) trp1::Gal-ISCeI-TRP1 Rad51-sfGFP* | Fig. 5D, 7E | [7] | W303 |
| ECS3453 | *rad52-F316A* | Fig. 5D | | |
| ECS3455 | *rad52-F337A* | Fig. 5D | | |
| ECS3457 | *rad52-Y376A* | Fig. 5D | | |
| yAT3974 | *Srs2::HIS3* | Fig. 5D, 7E | [7] | |
| ECS3531 | *rad52-F316A srs2 ::HPH* | Fig. 5D | | |
| ECS3533 | *rad52-F337A srs2 ::HPH* | Fig. 5D | | |
| ECS3535 | *rad52-Y376A srs2 ::HPH* | Fig. 5D | | |
| ECS3790 | *rad51-T161R* | Fig. 7E | | |
| ECS4003 | *rad51-T161R srs2 ::HIS3* | Fig. 7E | | |
| YMV80 | *MAT***a** *ade1-100 ura3-52 leu2-3,112 lys5 hml::ADE1 mat::hisG hmr::ADE1 leu2-cs his4::NAT-leu2Δ5′ ade3::GAL::HO* | Fig. 5-Ex1A | J. Haber | YFP17 |
| EMY496 | *rad52-F316A* | Fig. 5-Ex1A | | |
| ECS3630 | *rad52-F337A* | Fig. 5-Ex1A | | |
| TLM207 | *Rad52-Y376A* | Fig. 5-Ex1A | [6] | |
| TLM264 | *srs2 ::KanMX* | Fig. 5-Ex1A | | |
| ECS3623 | *srs2 ::KanMX rad52-F316A* | Fig. 5-Ex1A | | |
| ECS3626 | *srs2 ::KanMX rad52-F337A* | Fig. 5-Ex1A | | |
| TLM268 | *Srs2 ::LEU2 rad52-Y376A* | Fig. 5-Ex1A | [6] | |
| EMY493 | *rad52-F316A-6His-3FLAG-KANMX* | Fig. 5C, 5-Ext1B | | |
| ECS3402 | *rad52-F337A-6His-3FLAG-KANMX* | Fig. 5C, 5-Ext1B | | |
| ECS3220 | *RAD52-6His-3FLAG-KANMX srs2::HPH* | Fig. 5C, 5-Ext1B, 7D | | |
| ECS3404 | *rad52-F316A-6His-3FLAG-KANMX srs2::HPH* | Fig. 5C, 5-Ext1B | | |
| ECS3406 | *rad52-F337A-6His-3FLAG-KANMX srs2::HPH* | Fig. 5C, 5-Ext1B | | |
| ECS3785 | *RAD52-6His-3FLAG-KANMX rad51-T161R* | Fig. 7D | | |
| ECS3786 | *RAD52-6His-3FLAG-KANMX rad51-T161R srs2::HPH* | Fig. 7D | | |

| | | | | |
|---|---|---|---|---|
| ECS3222 | *rad52-Y376A-6His-3FLAG-KANMX srs2::HPH* | Fig 5C | [6] | |
| ECS3210 | *rad52-Y376A-6His-3FLAG-KANMX* | Fig 5C | [6] | |
| CTY10-5d | *MAT*a *ade2 trp1-901 leu2-3,112 his3-200 gal4 gal80 - URA3::lexAop-lacZ* | Fig. 6B, 6-Ex1A | S. Fields | CTY10-5d |
| TLM284 | *Rad51::KANMX* | Fig. 6B 6-Ex1A | | |

Strains are grouped according to their background, indicated on the first line of each group. Only differences from this genotype are noted subsequently.
